# Membrane Ca^2+^ permeability and IP3R2 dependent Ca^2+^-induced Ca^2+^ release are essential for astrocytic intracellular Ca^2+^ elevation upon neuronal stimulation at the mouse hippocampal CA3 - CA1 excitatory synapses

**DOI:** 10.1101/2020.10.19.345579

**Authors:** Jarand B. Hjukse, Gry Fluge Vindedal, Rolf Sprengel, Vidar Jensen, Erlend A. Nagelhus, Wannan Tang

**Author notes:** Deceased (Erlend A. Nagelhus).

## Abstract

Astrocytes are intricately involved in the activity of neural circuits, however, their basic physiology of interacting with neurons remains controversial. Using dual-indicator two-photon imaging of neurons and astrocytes during stimulations of hippocampal CA3 - CA1 Schaffer collateral (Scc) excitatory synapses, we report that under physiological conditions, the increased glutamate released from the higher frequency stimulation of neurons can accelerate local astrocytic Ca^2+^ levels. As consequences of extracellular glutamate clearance and maintaining of astrocytic intracellular Na^+^ homeostasis, the increase of astrocytic membrane Ca^2+^ permeability via Na^+^/Ca^2+^ exchanger (NCX) reverse mode is the primary reason of eliciting astrocytic intracellular Ca^2+^ elevation upon neuronal stimulation. This Ca^2+^-induced Ca^2+^ release is dependent on inositol triphosphate receptor type 2 (IP3R2). In addition, ATP released from Scc excitatory synapses can contribute to this molecular mechanism of Ca^2+^-induced Ca^2+^ release in astrocytes.

## Introduction

Modern neuroimaging technology has revolutionized our knowledge on the function of astrocytes, the predominant glial cell type in the brain. However, mechanisms underlying neuronal stimulation-evoked astrocytic Ca^2+^ signals are debated and vary between species, brain regions, developmental stages and astrocytic subcellular compartments (Saab et al. 2012; Doengi et al. 2009; Droste et al. 2017; Zhang et al. 2019; Santello, Toni, and Volterra 2019; Tang et al. 2015; Deitmer and Rose 2010). As important contributors to elicit astrocytic Ca^2+^ responses upon neuronal stimulation, various neurotransmitters, ion channels, G-protein coupled receptors, and neurotransmitter transporters embedded in the astrocytic membrane are currently involved in this controversy (Bazargani and Attwell 2016; Santello, Toni, and Volterra 2019; Deitmer and Rose 2010).

The basal level of astrocytic intracellular Ca^2+^ at the resting state is tightly regulated at low concentrations (about 100 nM or below), which is necessary for maintaining astrocytic physiological function (Deitmer and Rose 2010; King et al. 2020; Shigetomi et al. 2010). Unlike neurons, few pathways allowing extracellular Ca^2+^ entry are present in the astrocytic membrane, including receptors, channels, exchangers and pumps. This leads to a highly restricted Ca^2+^ permeability of the astrocytic membrane (Bazargani and Attwell 2016). However, whether the neuronal stimulation-evoked astrocytic Ca^2+^ increases are associated with relaxation of this limited astrocytic membrane Ca^2+^ permeability has remained unclear.

## Results and Discussion

At hippocampal CA3 - CA1 Schaffer collateral/commissural fibers (Scc), glutamate is the main neurotransmitter released at excitatory synapses. The glutamate release contributes to the increase of astrocytic intracellular Ca^2+^ as shown in our previous study (Tang et al. 2015). Thus, we first quantified the amount of glutamate released from the Scc synapses with three incremental stimulation protocols (20 Hz for 1 sec, 20 Hz for 10 sec and theta burst with 5 trains at 100 Hz repeated 5 times every 200 msec). The glutamate released from respective neuronal stimulation was measured by a genetically encoded fluorescent glutamate sensor iGluSnFR under *hSYN* promoter expressed with a rAAV vector (Marvin et al. 2013). Stimulation of Scc yielded a sharp increase in iGluSnFR fluorescence throughout the *stratum radiatum* (Figure 1A and 1B), albeit with lower amplitudes distant to the stimulating electrode (Figure 1B – 1D), showing iGluSnFR is capable of detecting the extracellular glutamate increase (Figure 1E – 1G and 1H **left**). Even during the high frequency theta burst stimulation, iGluSnFR transients did accompany every stimulation train, with a recovery time of 40 – 50 msec (Figure 1G and 1H **right**). Intriguingly, the theta burst protocol evoked an iGluSnFR fluorescence increase of significant higher amplitude than other two protocols (Figure 1H **left**), demonstrating a greater amount of glutamate release upon higher level of neuronal stimulation.

**Figure 1.**
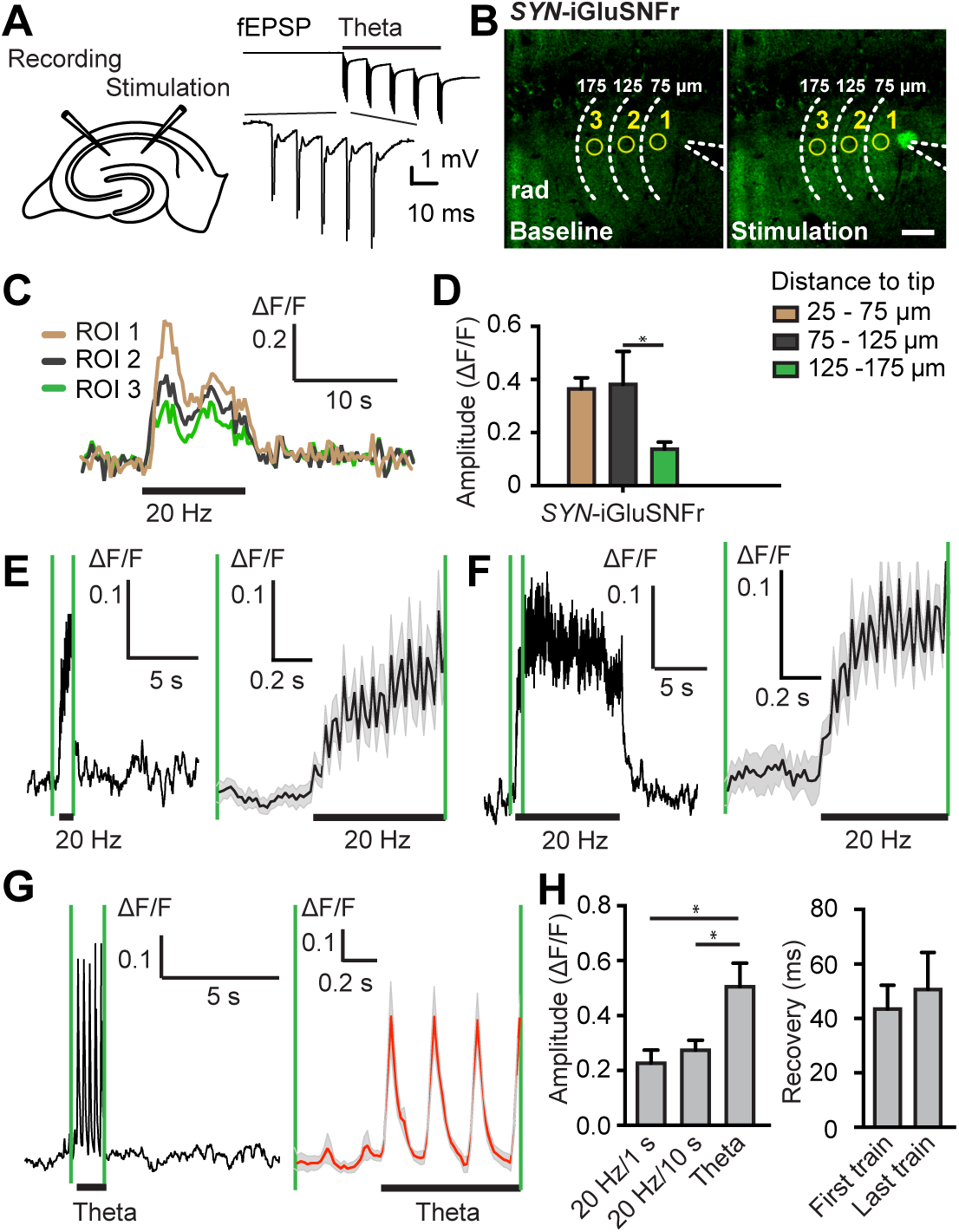
Neuronal stimulation-evoked elevation of extracellular glutamate levels. (A) Left, schematic drawing of the electrode placement in the hippocampal acute slice; right, representative trace of fEPSP (expanded in bottom trace) recorded during theta burst stimulations. (B) Images of *SYN*-iGluSnFr fluorescence in adult mouse hippocampus before (baseline) and after 20 Hz for 10 sec Scc stimulation. Dashed lines indicate position of the stimulation electrode and distance to the electrode tip. Scale bar, 10 µm; rad, *stratum radiatum*. (C) Representative traces of three regions of interest (ROIs 1-3) indicated in the images of (B). Black bar indicates 20 Hz for 10 sec Scc stimulation. (D) Amplitude of stimulation-evoked (20 Hz, 10 sec) extracellular iGluSnFr fluorescence at different distances from the stimulation electrode (P = 0.0059, n_25-75µm_ = 7, n_75-125µm_ = 7, n_125-175µm_ = 6). (E) – (G) Representative traces of *SYN*-iGluSnFr fluorescence during neuronal stimulation (indicated with bar) at 20 Hz for 1 sec (E), at 20 Hz for 10 sec (F) and theta burst (G). Traces between the green vertical lines are expanded at the right side. Grey shades indicate the range of standard error mean (SEM). (H) Left, amplitudes of stimulation-evoked extracellular iGluSNFr fluorescence (P = 0.0059, F_Genotype_ (2, 26) = 6.285, n_20Hz/1s_ = 9, _n20Hz/10s_ = 11, n_Theta_ = 9). Right, iGluSnFr fluorescence recovery rate of first and last train during theta burst stimulation (P = 0.8750, n = 8). Asterisk in (D) and (H) indicates values that differ significantly from each other.

To further investigate the dynamics of astrocytic intracellular Ca^2+^ increases in relation to the amount of glutamate released from the Scc synapses, we used wild-type mice transduced with a mixture of rAAV-h*SYN*-jRGECO1a (Dana et al. 2016) and rAAV-*GFAP-*GCaMP6f (Enger et al. 2015) to simultaneously reveal neuronal and astrocytic Ca^2+^ signals, respectively. Both 20 Hz for 1sec and theta burst stimulation of Scc (Figure 2A) evoked distinct increases in neuronal jRGECO1a (red) and astrocytic GCaMP6f (green) fluorescence (Figure 2B and 2C). In wild-type mice, the relative latency of astrocytic Ca^2+^ increases in all compartments occurred about 2.5 sec after the neuronal Ca^2+^ rise with 20 Hz for 1 sec stimulation (Figure 2D). However, a significant shorter relative latency in astrocytic somata and processes (about 1.7 sec) was detected by using theta burst stimulation (Figure 2D), indicating that a higher frequency stimulation resulted in a faster onset of astrocytic Ca^2+^ increases. Thus, together with the above iGluSnFR experiment, we show that the greater glutamate released from the higher frequency stimulation of neurons accelerated the local astrocytic Ca^2+^ increases.

**Figure 2.**
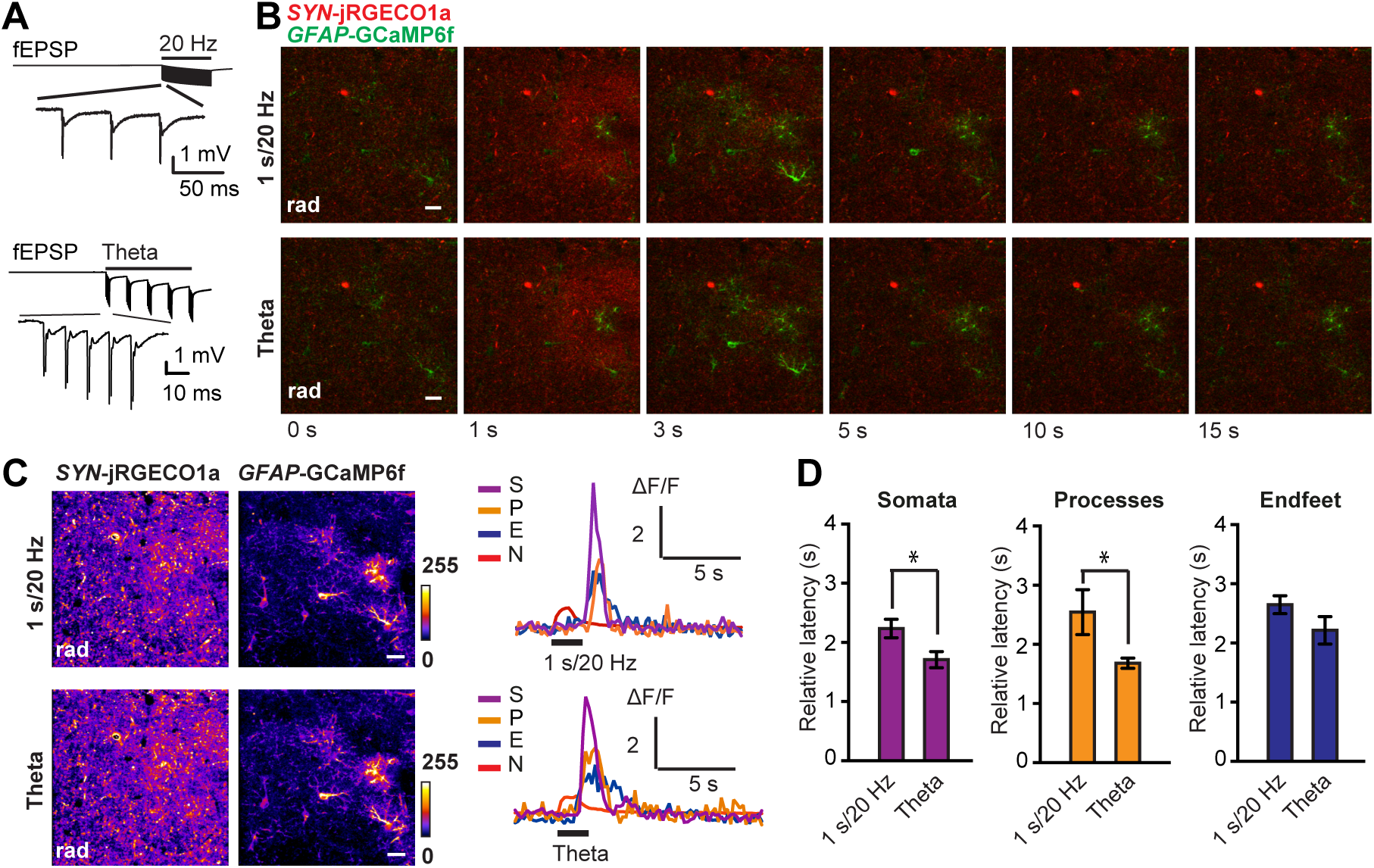
Dual-indicator two-photon imaging of Ca^2+^ signals in acute wild-type mouse hippocampal slices transduced with rAAV-*SYN*-jRGECO1a and rAAV-*GFAP*-GCaMP6f with two stimulation protocols. (A) Representative trace of fEPSP (expanded in bottom trace) recorded during 20 Hz for 1 sec (top) and theta burst (bottom) stimulation. (B) Time series of *GFAP*-GCaMP6f (green) and *SYN*-jRGECO1a (red) fluorescence images with two stimulation protocols of the Scc. (C) Left, standard deviation images of jRGECO1a and GCaMP6f fluorescence intensities from a time series recording with above mentioned two stimulation protocols; right, representative traces of the neuronal (N) and astrocytic Ca^2+^ signals with two stimulation protocols. S, astrocytic somata; P, astrocytic process. Black bar indicates Scc stimulation. (D) Bar diagram showing the latency between the onset of the neuronal jRGECO1a fluorescence transient and the one of astrocytic GCaMP6f fluorescence transients in somata (left), processes (middle) and endfeet (right) with two Scc stimulation protocols (P_somata_ = 0.035, n = 27; P_processes_ = 0.0023, n = 79; P_endfeet_ = 0.082, n = 16). Asterisks indicate values that differ significantly from each other. Scale bar, 10 µm, rad, *stratum radiatum*.

Glutamate transporters (GLT/GLAST) in astrocytic membranes are responsible for the uptake of neuronal released glutamate from the extracellular space (ECS). The entry of the glutamate via the transporter is accompanied by the elevation of intracellular Na^+^ concentration in astrocytes, which subsequently can activate the reverse mode of astrocytic Na^+^/Ca^2+^ exchanger (NCX) to export Na^+^ from astrocytes and to import Ca^2+^ into astrocytes (Deitmer and Rose 2010; Santello, Toni, and Volterra 2019). When we blocked the glutamate transporter with its specific inhibitor DL-threo-β-Benzyloxyaspartic acid (DL-TBOA), a severe reduction of neuronal field excitatory postsynaptic potential (fEPSP) about 75% (Supplementary Figure 1A) was observed, and astrocytic Ca^2+^ increases upon Scc stimulation (20 Hz for 10 sec) showed a significant reduction in processes (Supplementary Figure 1B). Although the rise time both in somata and processes was shortened, and the response duration in the somata was prolonged, astrocytic Ca^2+^ increases still remained high (Supplementary Figure 1B), demonstrating both glutamate transporters and other potential triggering mechanisms are simultaneously contributing to the rise of astrocytic Ca^2+^ signals.

As the uptake of glutamate in astrocytes could lead to a direction reverse of NCX, a potent and selective inhibitor of NCX reverse mode, KB-R7943, was applied to block the extracellular Ca^2+^ entry through NCX. We asked, whether this indirect pathway of increasing astrocytic Ca^2+^ membrane permeability through NCX is essential for triggering astrocytic Ca^2+^ increases. Surprisingly, with comparable neuronal stimulation strength revealed by the jRGECO1a signals before and after bath application of KB-R7943, astrocytic Ca^2+^ increases were almost completely abolished (Figure 3A – C and 1G). This suggested that a Ca^2+^-induced Ca^2+^ release mechanism involving the import of extracellular Ca^2+^ by the the reverse mode of NCX is essential for initiating the internal release of Ca^2+^ from ER stores. In addition, the astrocytic Ca^2+^ increases are abolished in *Itpr2*^−/−^ mice (Supplementary Figure 2), suggesting this NCX-mediated Ca^2+^-induced Ca^2+^ release is dependent on inositol triphosphate receptor type 2 (IP3R2).

**Figure 3.**
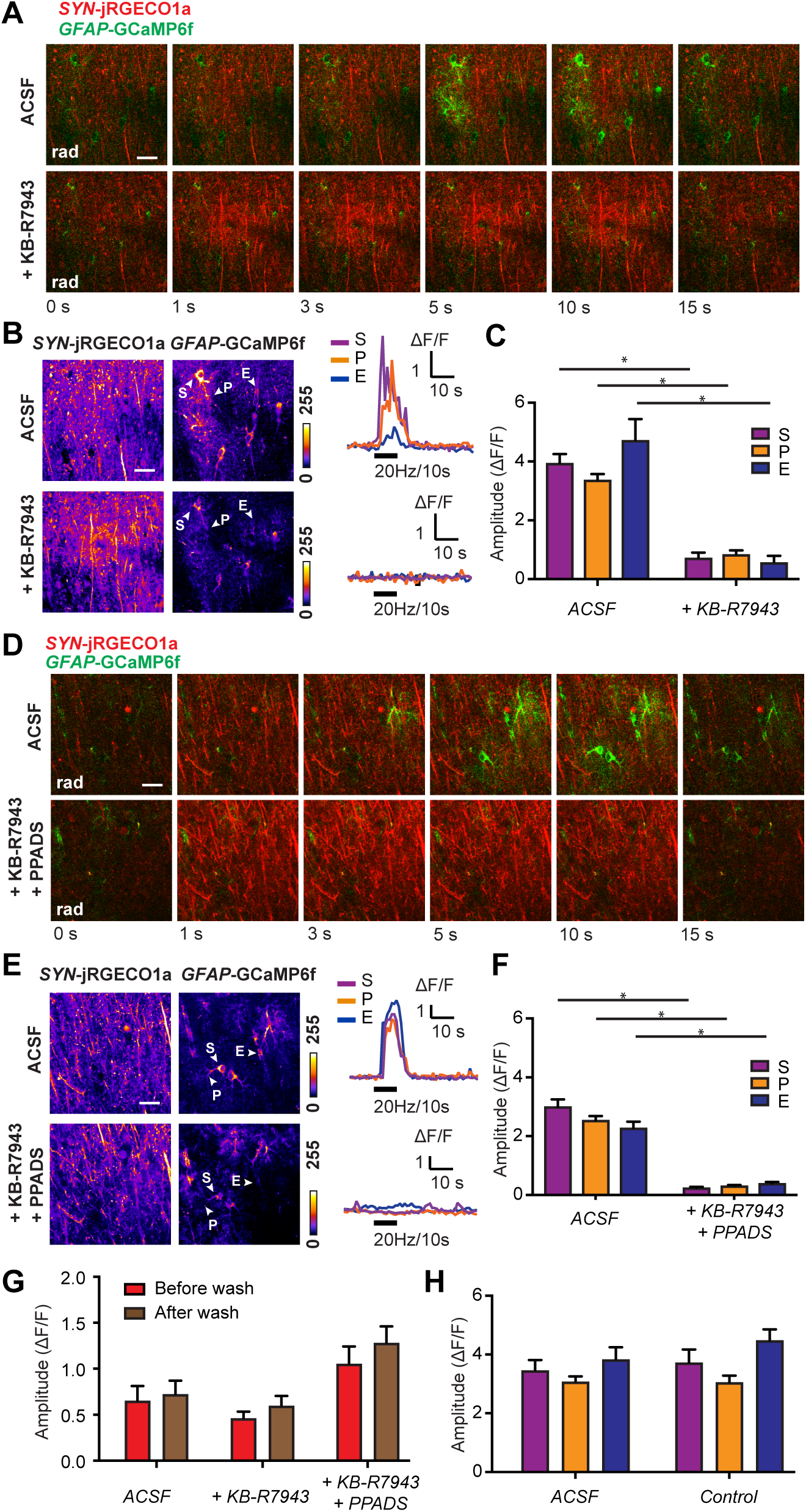
Increased astrocytic membrane Ca^2+^ permeability via NCX and P2XRs is essential for triggering neuronal-evoked astrocytic Ca^2+^ responses. (A) Time series of *GFAP*-GCaMP6f (green) and *SYN*-jRGECO1a (red) fluorescence images from wild-type adult mice Scc stimulated at 20 Hz for 10 sec with only ACSF and with KB-R7943 (20 μM) in ACSF. (B) Left, standard deviation images of jRGECO1a and GCaMP6f fluorescence intensities from above mentioned time series in a; right, traces of the astrocytic Ca^2+^ signals from the regions indicated with arrowheads in the images to the left. (C) Bar diagrams showing the amplitude (ΔF/F) of the GCaMP6f fluorescence increase in astrocytic compartments stimulated at 20Hz for 10 sec with only ACSF and with KB-R7943 in ACSF (P_treatment_ < 0.0001, F_Genotype_ (1, 303) = 164.7; P_interaction_ = 0.0290, F_interaction_ (2, 303) = 3.583, n_S-ACSF_ = 39, n_S-KB-R7943_ = 39, n_P-ACSF_ = 102, n_P-KB-R7943_ = 103, n_E-ACSF_ = 13, n_E-KB-R7943_ = 13). (D) Time series of *GFAP*-GCaMP6f (green) and *SYN*-jRGECO1a (red) fluorescence images from wild-type adult mice Scc stimulated at 20 Hz for 10 sec with only ACSF and with both KB-R7943 (20 μM) and PPADS (100 μM) in ACSF. (E) Left, standard deviation images of jRGECO1a and GCaMP6f fluorescence intensities from above mentioned time series in (D); right, traces of the astrocytic Ca^2+^ signals from the regions indicated with arrowheads in the images to the left. (F) Bar diagrams showing the amplitude (ΔF/F) of the GCaMP6f fluorescence increase in astrocytic compartments stimulated at 20Hz for 10 sec with only ACSF and with both KB-R7943 and PPADS in ACSF (P_treatment_ < 0.0001, F_Genotype_ (1, 292) = 351.4; P_interaction_ = 0.0306, F_interaction_ (2, 292) = 3.528, n_S-ACSF_ = 34, n_S-KB-R7943-PPADS_ = 31, n_P-ACSF_ = 82, n_P-KB-R7943-PPADS_ = 77, n_E-ACSF_ = 37, n_E-KB-R7943-PPADS_ = 37). (G) Bar diagrams showing with 20 Hz for 10 sec stimulation, the amplitude (ΔF/F) of jRGECO1a signals in neurons with only ACSF (P = 0.8750, n = 4), with KB-R7943 (P = 0.1289, n = 9), and with both KB-R7943 and PPADS (P = 0.0625, n = 6) blockage. (H) Bar diagrams showing with 20 Hz for 10 sec stimulation, the amplitude (ΔF/F) of astrocytic GCaMP6f signals in astrocytic compartments 1hour after ACSF wash (as the control for drug wash-in step, P_interaction_ = 0.5716, F_interaction_ (2, 242) = 0.5607, P_treatment_ = 0.2662, F_treatment_ (1, 242) = 1.242, n_S-ACSF_ = 30, n_S-Control_ = 34, n_P-ACSF_ = 68, n_P-Control_ = 72, n_E-ACSF_ = 22, n_E-KB-Control_ = 22). Asterisk in (C) and (F) indicates values that differ significantly from each other. Scale bar, 10 µm. rad, *stratum radiatum*; S, somata; P, processes; E, endfeet. Black bar indicates the electrical Scc stimulation.

However, when we repeated the same protocol with a higher neuronal stimulation strength, astrocytic Ca^2+^ increases could be measured despite the presence of KB-R7943. Since ATP is known as another important neurotransmitter released at excitatory synapses (Deitmer and Rose 2010; Bazargani and Attwell 2016), the binding of ATP to P2X receptor channels (P2XRs) could potentially lead to extracellular Ca^2+^ entry, thereby increasing the astrocytic Ca^2+^ membrane permeability. Therefore, we next performed experiments blocking both NCX reverse mode and P2XRs simultaneously with KB-R7943 and Pyridoxalphosphate-6-azophenyl-2’,4’-disulfonic acid tetrasodium salt (PPADS), respectively. During this double blockage, the Ca^2+^ increases in all microdomains of astrocytes were completely suppressed (Figure 3D – 3F) despite of a 2-fold increase in neuronal stimulation strength measured by the jRGECO1a fluorescence (Figure 3G). As expected, washing in of just artificial cerebrospinal fluid (ACSF) as controls did not change levels of astrocytic Ca^2+^ elevation (Figure 3H). These results identify extracellular Ca^2+^ influxes through NCX and P2XRs on the astrocytic membrane as primary initiators for the Ca^2+^ elevation in astrocytes. Thus, the astrocytic membrane Ca^2+^ permeability and the IP3R2 dependent Ca^2+^-induced Ca^2+^ release mechanism via NCX and P2XRs are essential for triggering the neuronal stimulation-evoked Ca^2+^ increase in astrocytes at the hippocampal CA3-CA1 Scc excitatory synapses of adult mice.

Here we confirmed that the neuronal released glutamate and ATP can trigger astrocytic Ca^2+^ increases at the hippocampal Scc, as suggested in our previous study (Tang et al. 2015). However, different to the pharmacological blockade of mGluRs and unspecific purinergic receptors that lead only to a partial reduction of Ca^2+^ elevation (Tang et al. 2015), we now discovered that the extracellular Ca^2+^ entry through NCX and P2XRs governs the astrocytic intracellular Ca^2+^ elevation with an IP3R2 dependent Ca^2+^-induced Ca^2+^ release mechanism. The astrocytic membrane Ca^2+^ permeability modulated by molecules in the extracellular space (ECS) plays the essential role on neuronal stimulation-evoked astrocytic activation.

In other brain regions, such as olfactory bulb, retina and cerebellum, studies have shown that extracellular Ca^2+^ entry via Ca^2+^ permeable AMPA receptors or GABA transporter as important causes for inducing astrocytic Ca^2+^ elevation upon local neuronal stimulation (Doengi et al. 2009; Saab et al. 2012; Zhang et al. 2019), supporting and generalizing our conclusion on the importance of astrocytic membrane Ca^2+^ permeability. Our results are pointing out that the change of the astrocytic membrane Ca^2+^ permeability is the driving force of the Ca^2+^ elevation in astrocytes when they are activated by local neurons (Figure 4), which is distinct from other (optional neuromodulatory) astrocytic activation pathways that involve mGluRs and G protein-coupled receptors.

**Figure 4.**
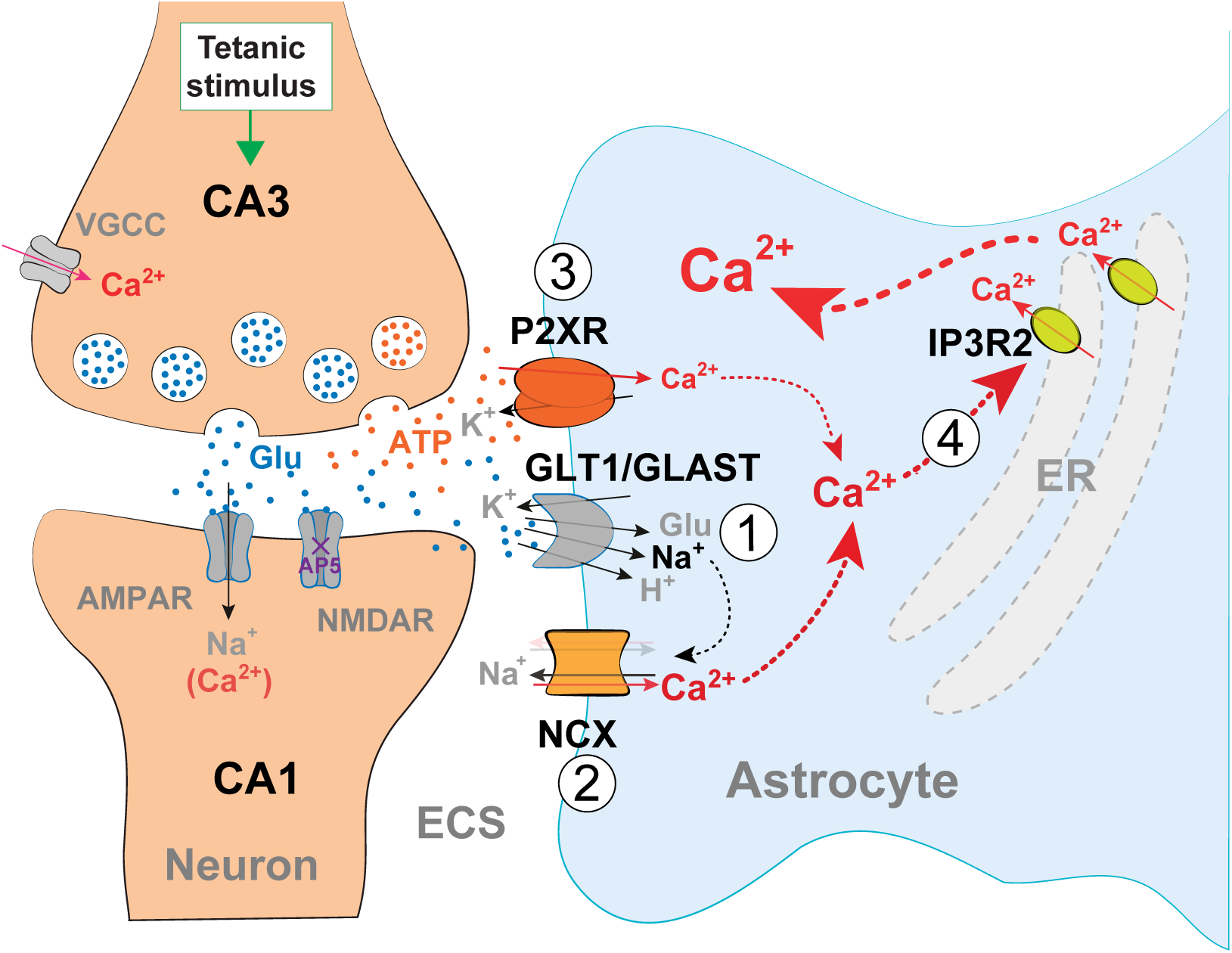
Astrocytic membrane Ca^2+^ permeability through NCX and P2XRs governs Ca^2+^-induced Ca^2+^ release through IP3R2 upon local neuronal stimulations. (**1**) The neuronal released glutamate (Glu) binds to glutamate transporters (GLT1/GLAST) on astrocytic membranes, and this could cause rapid Na^+^ influx to the astrocytes. (**2**) The elevated intracellular Na^+^ concentration in astrocytes could change the NCX from its forward mode (translucent arrows) to the reverse mode (solid arrows) and thus the import of Ca^2+^ into astrocytes. (**3**) In addition, the neuronal released ATP can promote directly Ca^2+^ influx at P2XR channels into astrocytes. (**4**) Together, the Ca^2+^ entering through NCX and P2XRs facilitates the Ca^2+^-induced Ca^2+^ release through IP3R2 on astrocytic ER.

## Materials and Methods

### Animals

Both male and female wild-type (C57BL6J, Janvier Labs) and *Itpr2*^−/−^ (*Itpr2*^*tm1*.*1Chen*^; MGI: 3640970)(Li et al. 2005) mice at least 6 weeks of age were housed with a 12-hour light/dark cycle (light on at 8 a.m.). *Itpr2*^−/−^ mice were backcrossed into a C57BL6J background for at least 15 generations. All experimental procedures were approved by the Norwegian Food Safety Authority (project number: FOTS 11255)

### Plasmid constructions and virus production

The DNA sequences for the genetically encoded fluorescent Ca^2+^ indicator GCaMP6f(Chen et al. 2013) and jRGECO1a(Dana et al. 2016) were first amplified by PCR from pGP-CMV-GCaMP6f and pGP-CMV-NES-jRGECO1a (Addgene) with 5’ BamHI and 3’ HindIII, and sub-cloned into the recombinant adeno-associated virus (rAAV) vector pAAV-6P-SEWB(Shevtsova et al. 2005) for generating pAAV-*SYN*-GCaMP6f and pAAV-*SYN*-jRGECO1a, respectively. The human glial fibrillary acidic protein (*GFAP*) promoter(Hirrlinger et al. 2009) was inserted with MluI and BamHI into pAAV-*SYN*-GCaMP6f construct for obtaining pAAV-*GFAP*-GCaMP6f. Plasmid pAAV-*SYN*-iGluSnFR(Marvin et al. 2013) was used to express the genetically encoded fluorescent glutamate indicator iGluSnFR. Serotype 2/1 rAAVs from pAAV-*GFAP*-GCaMP6f, pAAV-*SYN*-jRGECO1a and pAAV-*SYN*-iGluSnFR were produced(Tang et al. 2015) and purified by AVB Sepharose affinity chromatography(Smith, Levy, and Kotin 2009), following titration with real-time PCR (rAAV titer about 1.0 – 6.0 × 10^12^ viral genomes/mL, TaqMan Assay, Applied Biosystems). For hippocampal rAAV-transduction of both astrocytes and neurons, rAAV-*GFAP*-GCaMP6f and rAAV-*SYN*-jRGECO1a were mix 1:1.

### Surgical procedures and virus transduction

Viruses were stereotactically and bilaterally injected into the brains of deeply anesthetized (mixture of zolazepam (188 mg/kg body weight), tiletamine (188 mg/kg body weight), xylazine (4.5 mg/kg body weight) and fentanyl (26 µg/kg body weight)) 6 to 8-week-old C57BL6/J and *Itpr2*^−/−^ mice (Janvier Labs) as described(Tang et al. 2015). For transduction in adult mouse hippocampi, stereotactic coordinates relative to Bregma were: anteroposterior -2.0 mm, lateral ±1.5 mm. During injection, about 0.3 µl of purified rAAVs in total were delivered into each hippocampus with 1.5 mm in depth. All procedures were performed according to the guidelines of the local animal use and care committees.

### Electrophysiology and *ex vivo* two-photon Ca^2+^ imaging

Experiments were performed on acute hippocampal slices prepared from adult mice 2 to 6 weeks after rAAV transduction. The acute hippocampal slices were prepared as described(Tang et al. 2015) and kept at 30±1°C. Two glass electrodes filled with ACSF and positioned 100-150 µm away from each other in CA1 *stratum radiatum* served as stimulation and recording electrodes (fEPSP monitoring), respectively. In iGluSnFR experiments, DL-2-Amino-5-phosphonopentanoic acid (APV, 50 μM, Tocris) was added to the ACSF to avoid the unintended N-methyl-D-aspartate receptor (NMDAR) dependent plasticity. In order to stimulate the approximate same number of axons in the iGluSnFR experiment, we adjusted the stimulation strength so that we could record a prevolley of 1.0 mV in amplitude. In the GCaMP6f experiment we adjusted the stimulation strength to just below thresholds for eliciting a population spike on the third EPSP in a triple stimulation protocol (3 stimulation pulses with 20 ms interstimulus interval). In some experiments chemical blockers DL-TBOA (100 μm, Tocris), KB-R7943 (20 μM, Tocris), and PPADS (100 μM, Tocris) were added into ACSF. Recordings were done both before exposure to the drugs and 60 minutes post-exposure.

Stimulation in trains (20 Hz for 10 s, 20 Hz for 1 s, or theta burst (5 stimulation trains at 100 Hz repeated 5 times every 200 ms) were selectively applied during experiments. The GCaMP6f, jRGECO1a and iGluSnFR fluorescence were recorded by a two-photon laser scanning microscope (model “Ultima”, Prairie Technologies) with a “XLPLN 25×WMP” 1.05NA water-immersion objective (Olympus, Tokyo, Japan) at 900-910 nm (for imaging GCaMP6f and iGluSnFR) or 980-1020 nm (for dual-indicator imaging) laser pulse using a “Chameleon Vision II” (Coherent, Santa Clara, CA, USA) laser. Two-photon imaging was performed with 30 s baseline followed by electrical stimulations. The recordings were done either with 1 to 5 Hz for low imaging sampling rate (512×512 px or 256×256 px), or 50 Hz for high imaging sampling rate (50×50 px, for iGluSnFR experiment).

### Imaging analysis

Time-series of fluorescence images were analyzed with custom-written MATLAB (R2011b, MathWorks, Inc.) scripts. Regions of interests (ROIs) defining astrocytic compartments were selected over somata, processes and endfeet according to typical astrocyte morphology from the GCaMP6f signal. ROIs over processes were chosen at least 5 micrometer away from the perimeter of the somata. For dual-indicator latency experiment, circular ROIs of 3-5 µm in diameter were placed for both channels to obtain GCaMP6f and jRGECO1a signals simultaneously from astrocytes and neuropils, respectively. In the iGluSnFR experiment, circular ROIs of 5 µm were placed. The relative change in fluorescence (ΔF/F) in each ROI, the individual traces and the histograms were all calculated and plotted by MATLAB (R2011b, MathWorks, Inc.) with custom written scripts. To identify fluorescent events, 3 times standard deviation (SD) of baseline values was used as thresholding.

### Statistical analysis

Statistical analysis was performed using Prism (Version 8.0.2 for Mac OSX, GraphPad Software). Full descriptions of statistical parameter were accessed with the original data before choosing the suitable analysis. Ordinary one-way ANOVA was used for amplitude comparison of different stimulation protocols of iGluSnFR. Kruskal-Wallis test was used for amplitudes with distances measurement of iGluSNF signals. Wilcoxon test was used for measurements of recovery rate of iGluSnFR fluorescence after first, last train during theta burst stimulation, and control experiments of neuronal stimulation strength before and after drug incubation. Paired t test was performed with the relative latency of dual-indictor measurement and DL-TBOA experiment. Two-way ANOVA with multiple comparisons test (either Sidak’s multiple comparisons test, or Tukey’s multiple comparisons test) was performed with KB-R7943 and KB-R7943 + PPADS experiments, ACSF wash control and *Itpr2*^−/−^ mice experiments.

## Acknowledgements

We dedicate this work to Prof. Erlend A. Nagelhus, who tragically died on 10^th^ January 2020. We thank Prof. Loren L. Looger for providing all genetically encoded sensors, and Johannes Helm for technical support on the two-photon microscope. This work was supported by the Research Council of Norway Grant Number 262552, the European Union’s Seventh Framework Programme for research, technological development and demonstration under grant no. 601055, and the Letten Foundation.

## Author Contributions

Conceptualization and experimental design: E.A.N., V.J., W.T.; experimental conduct and methodology: J.B.H., G.F.V., R.S., V.J., W.T.; writing: R.S., W.T.; funding: E.A.N., W.T.

---

**Supplementary Figure 1.**
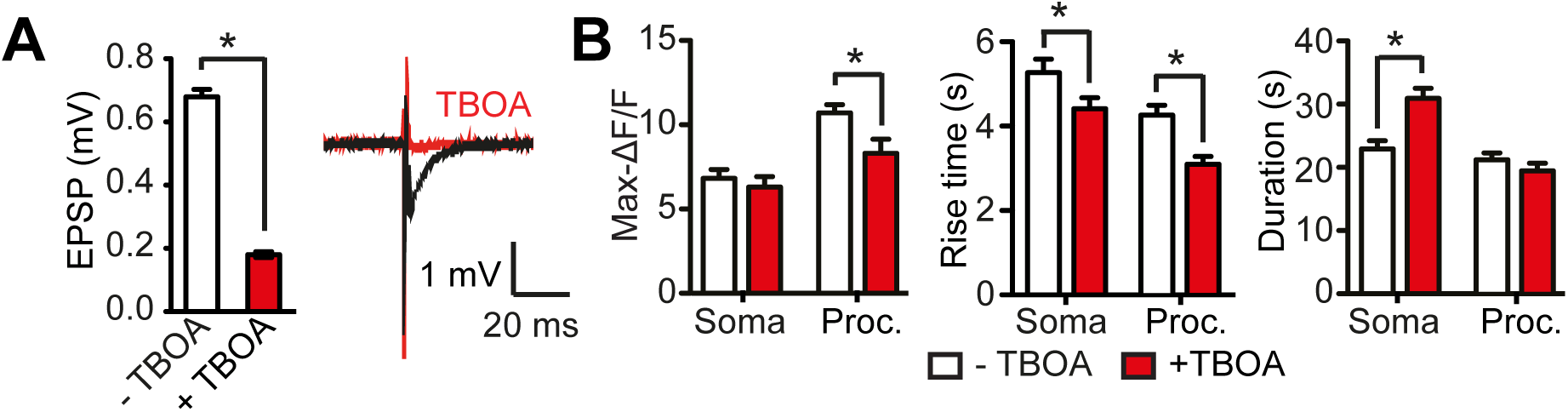
**(A)** Quantification of neuronal field EPSPs before and after DL-TBOA wash (left, P < 0.0001, n = 7) and representative traces (right). **(B)** After the DL-TBOA application, the maximum ΔF/F values dropped significantly in the processes (P = 0.0020, n = 60), rise time decreased in both soma and processes areas (P_soma_ = 0.0188, n = 41, P_processes_ < 0.0001, n= 60). The activation duration was increased in the soma after DL-TBOA wash in (P < 0.0001, n=60) whereas in the processes it was not significantly changed (P = 0.1498, n = 60).

**Supplementary Figure 2.**
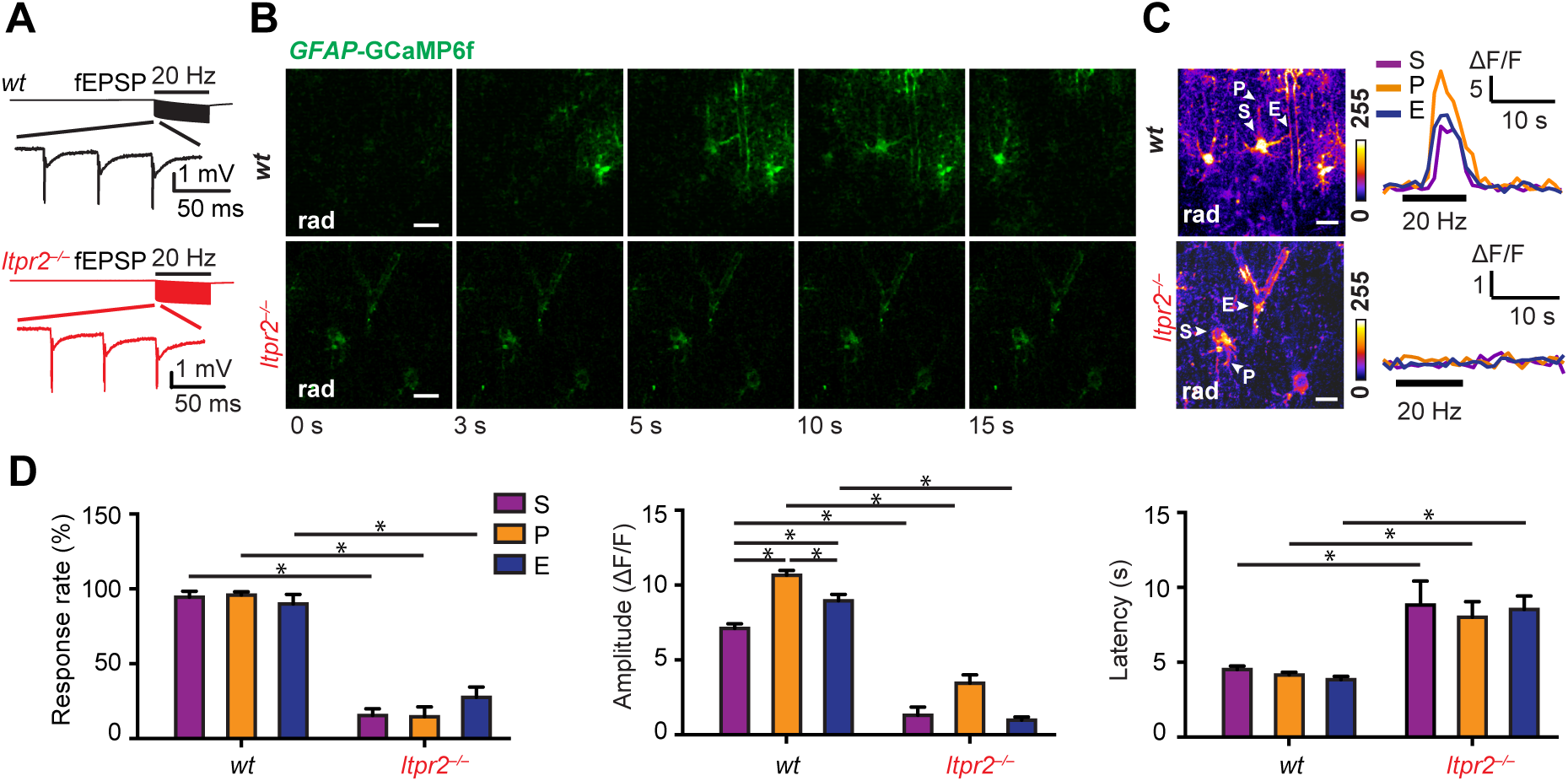
Neuronal stimulation-evoked astrocytic Ca^2+^ elevation is abolished by inositol triphosphate type 2 receptor (IP3R2) deletion revealed by *GFAP*-GCaMP6f fluorescence. **(A)** The fEPSPs (expanded in bottom trace) recorded during 20 Hz, 10 seconds stimulation in wild-type (*wt*, traces in black) and *Itpr2*^−/−^ (traces in red) mice. **(B)** Time series of *GFAP*-GCaMP6f fluorescence images from two genotypes (*wt* and *Itpr2*^−/−^) during the electrical stimulation (20 Hz, 10 s) of the Schaffer collateral/commissural fibres (Scc). Scale bar, 10 µm, rad, *stratum radiatum*. **(C)** Left, standard deviation images of *GFAP*-GCaMP6f fluorescence intensities during electrical stimulation (20 Hz, 10 s) in *wt* and *Itpr2*^−/−^ mice. Scale bar, 10 µm; rad, *stratum radiatum*; right, representative fluorescence traces from astrocytic compartments indicated in corresponding images on the left. S, somata; P, processes; E, endfeet. **(D)** Response rate (two-way ANOVA, P_Genotype_ < 0.0001, F_Genotype_ (1, 107) = 294.2; n_wt-S_ = 20, n_wt-P_ = 20, n_wt-E_ = 20, n_*Itpr2*–/– -C_ = 18, n_*Itpr2*–/– -P_ = 17, n_*Itpr2*–/– -E_ = 18), amplitude (two-way ANOVA, P_Genotype_ < 0.0001, F_Genotype_ (1, 453) = 139.4; P_Compartment_ = 0.0004, F_Compartment_ (2, 453) = 7.993; n_*wt*-S_ = 130, n_*wt*-P_ = 159, n_*wt*-E_ = 115, n_*Itpr2*–/– -C_ = 11, n_*Itpr2*–/– -P_ = 17, n_*Itpr2*–/– -E_ = 27), and latency (two-way ANOVA, P_Genotype_ < 0.0001, F_Genotype_ (1, 464) = 99.03; n_*wt*-S_ = 128, n_wt-P_ = 171, n_*wt*-E_ = 115, n_*Itpr2*–/– -C_ = 11, n_*Itpr2*–/– -P_ = 18, n_*Itpr2*–/– -E_ = 27) of neuronal stimulation-evoked (20 Hz, 10 s) GCaMP6f fluorescence responses in astrocytic somata (S), processes (P) and endfeet (E) in *wt* and *Itpr2*^−/−^ mice. Asterisks and lines indicate values that differ significantly from each other (response rate, Sidak’s multiple comparisons test, P < 0.05; amplitude, Tukey’s multiple comparisons test, P < 0.05; latency, Sidak’s multiple comparisons test, P < 0.05).

